# High-throughput dual bioreporter screening reveals distributed regulation of biofilm matrix components in *Staphylococcus aureus*

**DOI:** 10.64898/2026.08.04.742763

**Authors:** Jean-Sébastien Bourassa, Élizabeth Gaudreau, Jean-Philippe Côté, Pascale B. Beauregard

## Abstract

*Staphylococcus aureus* biofilm formation is a key factor enabling persistent infections. However, the lack of efficient high-throughput tools previously limited systematic study of its regulatory mechanisms. Here, we used high-efficiency transduction to construct two luminescent bioreporter libraries, each probing a distinct biofilm regulatory pathway. Derived from the Nebraska Transposon Mutant Library, these libraries enabled rapid, quantitative screening of biofilm-associated gene expression in a high-throughput format. Our screens revealed a surprising lack of overlap in the regulation of the two biofilm components investigated: adhesin synthesis and extracellular DNA production. However, we identified *mntR* as a key gene involved in the expression of both biofilm components and confirmed the previously reported role of *yjbH*. Cross-lineage validation showed that these regulators retain conserved significance across multiple *S. aureus* backgrounds, although their phenotypic effects varied across strains. Collectively, this work provides a versatile, high-throughput framework to dissect the regulatory networks underlying complex phenotypes in *S. aureus*.

**Importance:** Biofilm formation is a major contributor to the persistence and treatment failure of *Staphylococcus aureus* infections, yet its regulatory network remains incompletely understood. We developed a high-throughput bioreporter platform that enables genome-wide screening of biofilm-associated gene expression across nearly 2,000 transposon mutants. Using this approach, we show that key biofilm processes, adhesion and extracellular DNA release, are controlled by largely distinct regulatory networks, and we identify *mntR* as a previously unrecognized regulator shared by both pathways. Beyond these biological insights, our work provides a versatile and readily adaptable strategy for dissecting complex regulatory systems in *S. aureus* and other bacterial species.

## Introduction

*Staphylococcus aureus* is a major threat to human health due to its wide range of infections, some of which are associated with mortality rates as high as 20% (1). The increasing prevalence of methicillin-resistant *Staphylococcus aureus* (MRSA) limits the efficacy of first-line antibiotics, such as anti-staphylococcal penicillins and first-generation cephalosporins (2), thereby necessitating alternative therapies (3, 4). Although current antibiotics effectively kill the pathogen in vitro, treatment failure remains a significant clinical issue (4, 5). These failures can be attributed to several factors, including the development of small colony variants (SCVs) or the presence of a biofilm, in which the bacteria are embedded in a self-secreted extracellular matrix (6, 7).

The biofilm matrix confers several advantages to its inhabitants, including reduced antibiotic diffusion and increased resistance to immune clearance (8, 9). These factors can lead to incomplete pathogen elimination and, in severe cases, recurrent infections that may progress to a chronic, life-threatening condition (10). The matrix of MRSA biofilms mainly consists of Microbial Surface Components Recognizing Adhesive Matrix Molecules (MSCRAMMs) and extracellular DNA (eDNA) (11). One key MSCRAMM is the fibronectin-binding protein A (FnbA), which plays a crucial role in the initial adhesion phase of the biofilm. eDNA is primarily produced through programmed cell death, during which a subset of cells undergoes lysis due to a high murein hydrolase activity. This process is regulated by the CidA/LrgA, holin/anti-holin system. A decrease in either of these two biofilm components (MSCRAMMs or eDNA) reduces the ability of *S. aureus* to colonize the infection site and persist during infection (12, 13).

Despite their importance and the significant effort invested, studies of *S. aureus* biofilms remain hindered by technical challenges (14). A key limitation is the lack of real-time monitoring experiments and the necessity to use disruptive experiments to study this dynamic phenotype. In *Bacillus subtilis*, a widely used non-invasive technique for studying biofilms employs fluorescent bioreporters whose transcription is regulated by the same promoters that control extracellular matrix genes (15, 16). These bioreporters enable rapid, dynamic monitoring of biofilm formation and can be easily normalized to bacterial density. This approach has been used to quantify the impact of different compounds or interaction partners on *B. subtilis* biofilm production in real time (16, 17).

Here, we used the rapid quantitative readout of bioreporters to investigate biofilm formation in *S. aureus*. Two luminescent bioreporters, based on the *fnbA* and *cidA* transcriptional units (P*_fnbA_* and P*_cidA_*), were engineered to monitor the production of the two main biofilm matrix components. These reporters provide a non-destructive, real-time proxy for regulatory activity, enabling scalable screening of this complex phenotype.

This approach was then combined with the Nebraska Transposon Mutant Library (NTML), which comprises 1,920 strains with disruptions in most non-essential genes, to obtain a comprehensive view of biofilm regulatory determinants. Overall, our screen revealed limited overlap in the regulation of P*_fnbA_* and P*_cidA_*, suggesting that adhesin production and eDNA release are controlled by largely distinct regulatory pathways. Despite this divergence, we identified *yjbH* and the manganese-dependent transcriptional regulator *mntR* as key modulators of biofilm. Validation in alternative genetic backgrounds confirmed that YjbH and MntR are conserved regulatory nodes in *S. aureus* biofilm architecture, although their phenotypic effects vary across strains.

## Results

### Creation of P*_fnbA_* and P*_cidA_* reporter libraries

To monitor the dynamic production of the main biofilm matrix components, two biofilm bioreporters were constructed (Fig. 1A). The gene encoding fibronectin-binding protein A (*fnbA*) was used as a proxy for surface protein production, given its prominent role in early biofilm formation. To assess eDNA release, we monitored *cidA* expression, which encodes a holin central to this process. The promoters of these genes were cloned upstream of the luciferase operon engineered for Gram-positive bacteria (*luxABCDE*) to generate two transcriptional reporter plasmids (P*_fnbA_*-*luxABCDE* and P*_cidA_*-*luxABCDE*).

**Figure 1.**
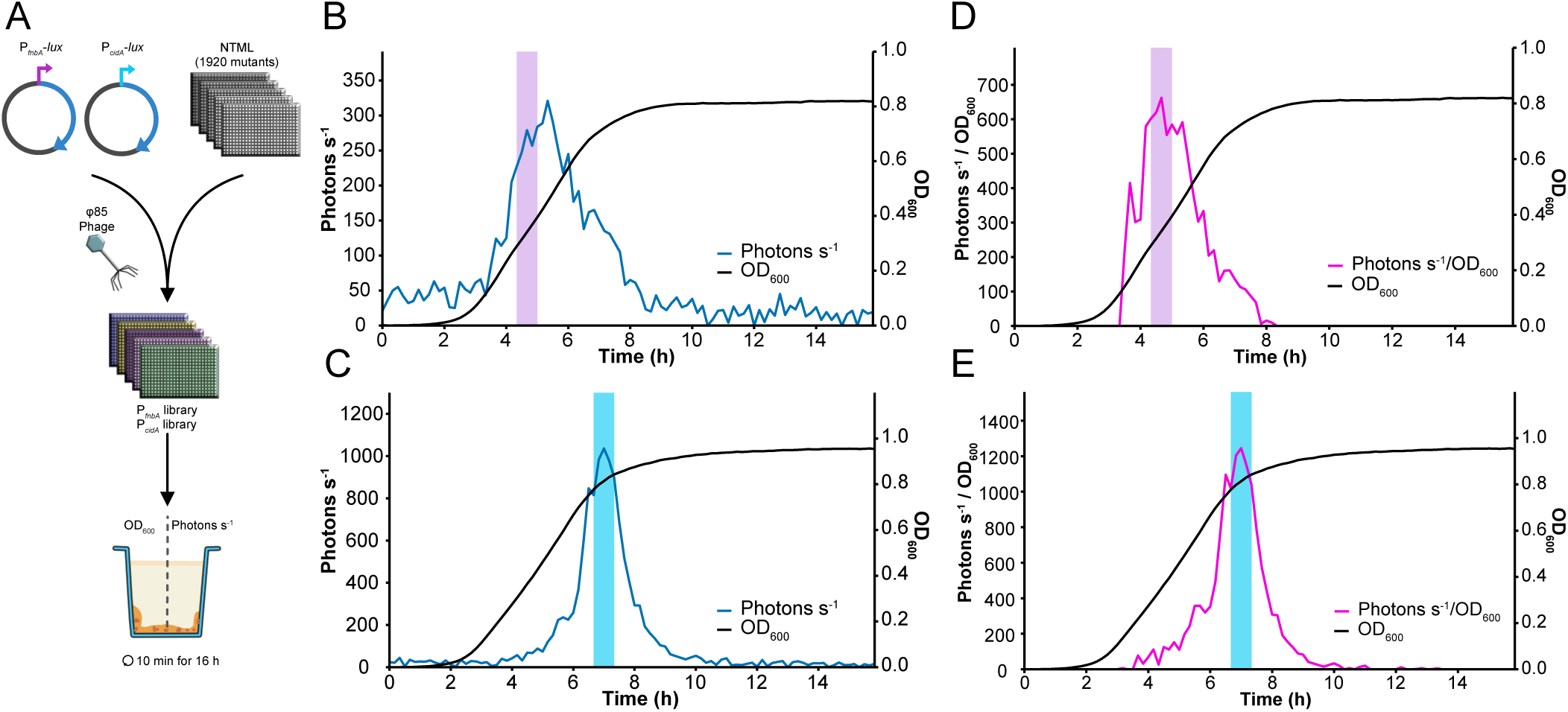
Overview of the high-throughput reporter library construction and screening workflow. (A) The main biofilm components were targeted by placing the *fnbA* promoter (MSCRAMMs) or the *cidA* promoter (eDNA) upstream of *luxABCDE* and then transducing the resulting constructs into the NTML to generate two reporter libraries. Raw reporter activity was measured from P*_fnbA_* (B) and P*_cidA_* (C) during biofilm growth. The P*_fnbA_* (D) and P*_cidA_* (E) curves were normalised bacterial density to identify the peak activity for further screening. Light pink (P*_fnbA_-lux*) and light cyan (P*_cidA_-lux*) rectangles represent the time point associated with the peak activity of each reporter (B-E).

*S. aureus* strain JE2 containing either plasmid was grown statically in liquid TSBg (Tryptic Soy Broth supplemented with 0,5% [w/v] glucose) to determine the expression kinetics of the reporters during biofilm formation. Although both FnbA and CidA are required for mature and robust biofilm formation by *S. aureus* JE2, they act at distinct stages of biofilm development, with FnbA involved in the initial adhesion step and CidA in biofilm maturation (18). Both reporters exhibited a sharp increase in signal, followed by a rapid decline, each occurring at different stages of bacterial growth. As expected, P*_fnbA_* activity increased during the early exponential phase (Fig. 1B). In contrast, P*_cidA_* activity peaked toward the end of the exponential phase (Fig. 1C). We then normalized these data to account for bacterial density (see method) and identified the peak activity of both reporters for further analysis (Fig. 1D, E). These results highlight that the luminescent bioreporters can dynamically monitor gene expression. We next validated that the bioreporter signals accurately reflect *fnbA* and *cidA* transcript levels by comparing their expression via RT-qPCR. As shown in Figure S1, the transcription levels of both P*_fnbA_*-*luxABCDE* and P*_cidA_*-*luxABCDE* were similar to those of the native genes in the WT background as well as in a mutant known to affect the transcription of the bioreporter (*saeR::tn* for *fnbA* and *cidR::tn* for *cidA*), thus, corroborating the reliability of the bioreporters.

To assess the impact of non-essential genes on biofilm formation, two libraries, carrying their respective bioreporter plasmid, were created by transducing the plasmids into each of the 1,920 NTML mutant strains. High-throughput transduction using phage ϕ85 was performed, and any remaining phages were carefully cleared by successive passages in TSB medium containing sodium citrate. Only the *menD::tn* strain could not be obtained with either bioreporter plasmid because of its intrinsic resistance to chloramphenicol, which was used for plasmid selection.

### P*_fnbA_* activity is highly impacted by energy deprivation

To assess the impact of the deleted genes on bioreporter activity, libraries were grown under static conditions in 384-well plates, and both growth (OD_600_) and reporter activity (Photons s^-1^) were monitored. As previously, we identified the peak activity and its related bacterial density for each strain (Fig. S2). Notably, examination of the P*_fnbA_* reporter library data first revealed two distinct subpopulations: one with a log_2_ relative activity ranging from 0 to 0.5, and another from −0.6 to −0.2 (Fig. 2A, Fig. S3). Whole-genome and plasmid sequencing of randomly selected strains from each population revealed identical plasmid sequences and no conserved genomic mutation that could account for this divergence. Plate analysis showed no plate pattern associated with these two populations. Further analysis indicated that part of the population showed a slight decrease in reporter activity, and they were no difference in growth between the two populations (Fig. S3B). These findings suggest that the subpopulations might arise from a specific intrinsic signalling pathway that regulates P*_fnbA_* expression.

**Figure 2.**
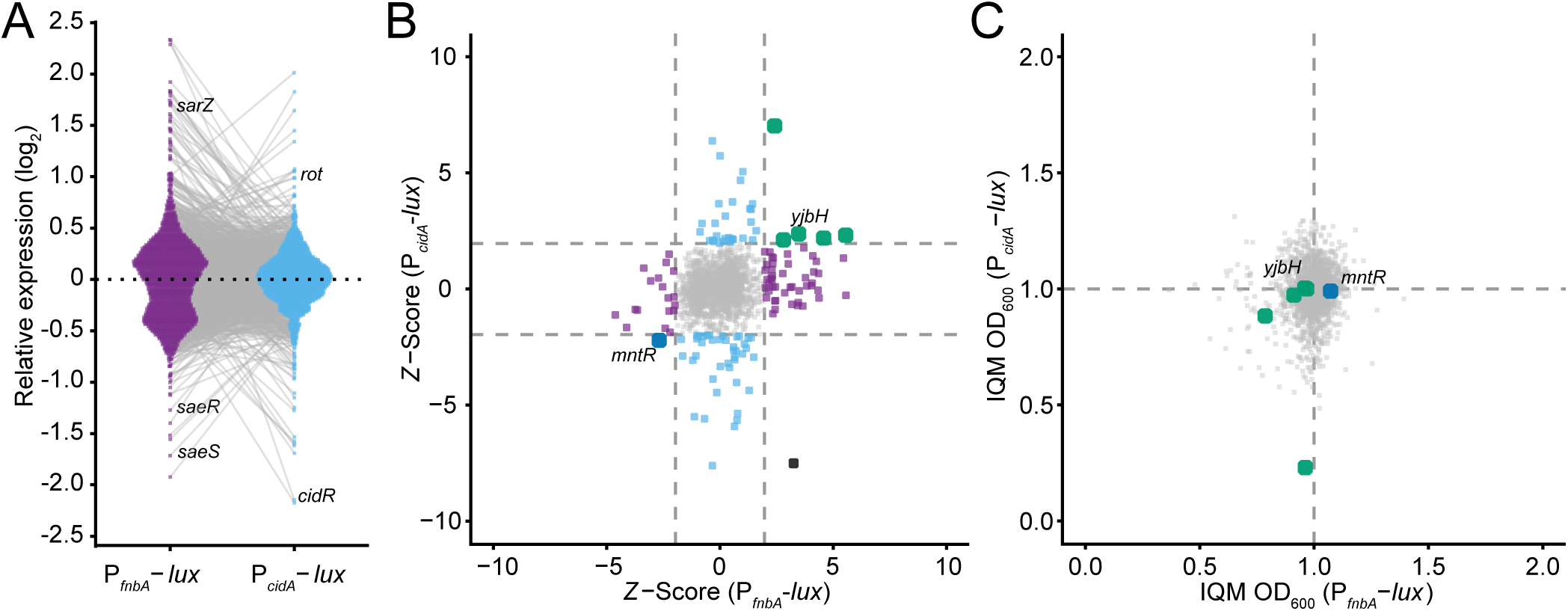
Genome-wide reporter screens identify distinct, limited sets of regulators that affect P*_fnbA_* and P*_cidA_* activity. (A) Distribution of the peak reporter activity across both libraries. (B) Comparison of reporter activity for each mutant present in both libraries, based on their respective z-scores. Applying a |z-score|>1.96 threshold identifies only a small set of regulators affecting both biofilm components. Mutants were classified according to their impact on each reporter. (C) Comparison of peak activity timing for both reporters shows no bias in peak detection when normalized using the IQM method. Purple dots represent the distribution of mutants within the P*_fnbA_*-*lux* library (A), or mutants significantly impacting the P*_fnbA_*-*lux* reporter only (B). Cyan dots represent the distribution of mutants within the P*_cidA_*-*lux* library (A), or mutants significantly impacting the P*_cidA_*-*lux* reporter only (B). Black dots represent the mutant that affects both reporters in different ways (B). Dark blue dots represent the mutant that negatively impacts both reporters (B-C), and green dots represent mutants that positively impact both reporters (B-C).

To identify mutants affecting bioreporter activity, we converted peak activity to a z-score to account for variation across the library (Fig. 2B). Mutants with |z-score| > 1.96 were considered to have a significant impact on bioreporter activity. Fifty-two mutants increased P*_fnbA_*, while 19 reduced its activity (Fig. 2). Consistent with its role as an activator of *fnbA* (11), the SaeRS two-component system emerged as a top hit, and its disruption reduced P*_fnbA_* activity (Table S1). Loss of *sarZ*, a member of the SarA regulator family, had the opposite effect, increasing P*_fnbA_* reporter activity (Table S1). The identification of these known regulators validates the robustness of the screening approach using the bioreporter library.

Hits identified from the library screen were subjected to Gene Ontology (GO) enrichment analysis using Biocyc to identify regulatory pathways associated with altered P*_fnbA_* activity. Mutants with increased P*_fnbA_* activity showed significant enrichment for GO terms related to energy metabolism and the DNA restriction-modification system (Fig. 3A). Although the restriction-modification system was the most significantly enriched category, its contribution may reflect effects on plasmid maintenance or copy number rather than a direct regulatory role in P*_fnbA_* expression.

**Figure 3.**
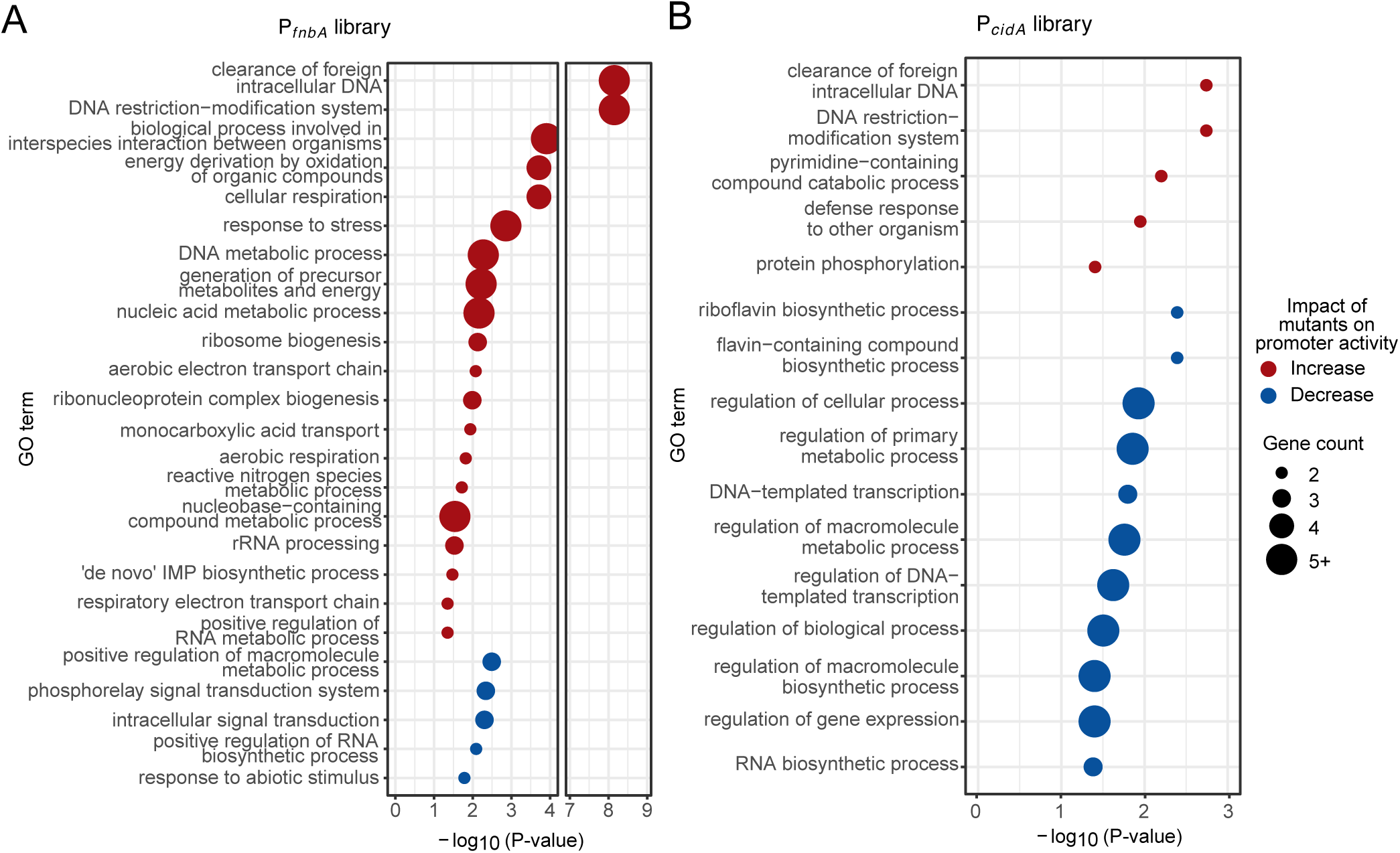
Functional enrichment analysis reveals pathways associated with altered reporter activity of P*_fnbA_* and P*_cidA_*. Statistical enrichment was assessed using a Fisher’s exact test without correction, applying a threshold of 0.05 and a minimum gene count of 2. Redundant or synonymous GO-terms were removed, keeping the more specific term if all previous genes were present. (A) Enriched GO-term among mutants that impact P*_fnbA_* activity. (B) Enriched GO-term among mutants that impact P*_cidA_* activity. GO-term enrichment in mutants that increase reporter activity is shown in red, and in mutants that decrease reporter activity in blue (A-B).

### Riboflavin biosynthesis genes are modestly enriched among hits that affect P*_cidA_*

In the P*_cidA_* reporter library, the screen identified 34 mutants that upregulated reporter activity and 43 that downregulated it (Fig. 2B). Notably, inactivation of *cidR*, the direct activator of *cidA,* abolished reporter activity (Table S1). In contrast, inactivation of the transcriptional regulator, *rot*, previously shown to limit biofilm formation (19), resulted in increased P*_cidA_* activity (Fig. 2A, Table S1). The identification of both mutants, previously characterized for their impact on biofilm formation, supports the reliability of this second library screen.

GO term enrichment analysis of the mutants identified in the P*_cidA_* reporter screen revealed enrichment of genes associated with the riboflavin biosynthetic process in mutants exhibiting reduced P*_cidA_* activity (Fig. 3B). Mutants with increased P*_cidA_* activity did not show clear enrichment of biological function (Fig. 3B).

### Distinct pathways with minimal overlap regulate P*_fnbA_* and P*_cidA_*

The screening of the two libraries highlighted both known and novel genes that may regulate extracellular matrix production. However, FnbA and CidA serve distinct roles within the biofilm and contribute to different phenotypes. To determine whether common regulatory pathways govern different aspects of biofilm formation in *S. aureus*, we compared the results of the two reporter screens. Only two GO terms, one associated with the restriction-modification system and a general term (regulation of biological process), were enriched in both reporter libraries (Fig. 3). We identified 141 mutants that affected the activity of at least one reporter. Of these, only six showed a similar impact on the two biofilm reporters (Fig. 2C, Table S1). We also identified one mutant, *recD2::tn*, that displayed opposing effects, up-regulating P*_fnbA_* while down-regulating P*_cidA_*. Five mutants (*hsdR::tn*, *hsdS2::tn*, *yjbH::tn*, *SAUSA300_0329::tn*, and *SAUSA300_0831::tn*) up-regulated the activity of both bioreporters, whereas *mntR::tn* led to a decrease in their activity. We evaluated the bacterial density at peak activity for these mutants to confirm that these hits are not biased toward higher or lower bacterial density (Fig. 2C). The top hit from the P*_cidA_* library is *SAUSA300_0831*, which also appears to allow earlier expression of P*_cidA_* than in the other mutants (Fig. 2B-C, S4, Table S1). Upon further analysis of this mutant, we confirmed that its deletion led to prompt signal activity, and that its expression (non-normalized) increased 3-fold relative to the WT (Fig. S4), supporting the validity of this hit rather than a bias in normalisation.

The mutants that affected both reporters similarly were then tested for their ability to adhere to fibronectin, the primary target of FnbA. Interestingly, the *yjbH::tn* mutant, which had higher P*_fnbA_* activity, showed increased in fibronectin adhesion (Fig. 4A). Inactivation of *mntR,* which reduced *fnbA* expression, led to a significant decrease in fibronectin binding. Complementation from a multicopy plasmid restored the adhesion ability of the *yjbH::tn* mutant to its WT level (Fig. 4A). Complementation of *mntR* increased the ability of this mutant to adhere to fibronectin, even to a level higher than the WT (Fig. 4A).

**Figure 4.**
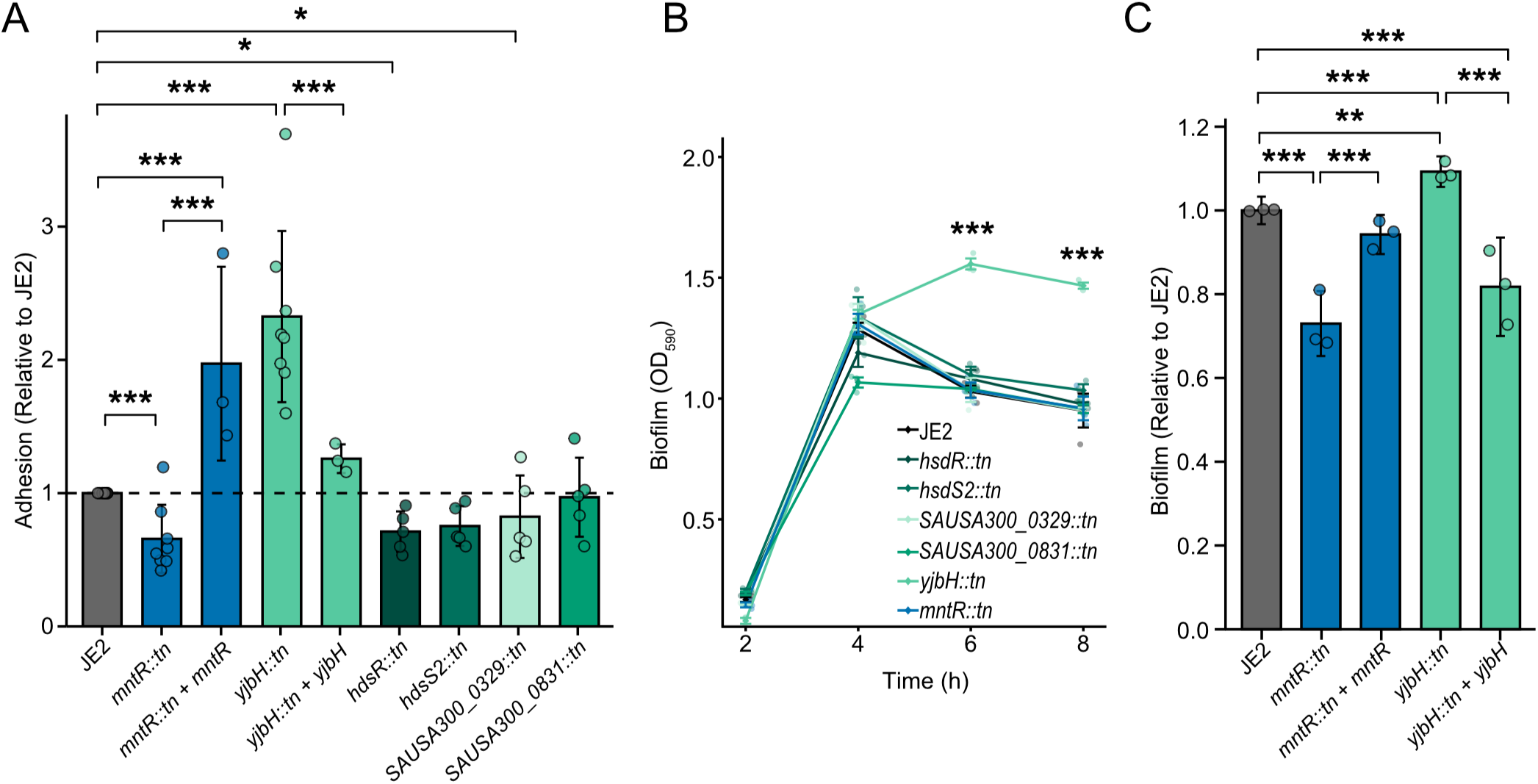
Validation of candidate mutants reveals altered adhesion to fibronectin and biofilm formation. (A) Adhesion of candidate mutants to fibronectin was measured to validate the phenotype identified with the P*_fnbA_* reporter. (B) Biofilm formation in TSBg was quantified by crystal violet staining over time. (C) Biofilm formation of complemented mutants was quantified using crystal violet staining after 8 h of growth in TSBg. Data are presented as adhesion (A) or biofilm (C) normalized to the WT for easier comparison between biological replicates. Data were analyzed using a linear (B) or linear mixed-effects model (A, C) followed by a Type III ANOVA and Dunnett-adjusted post hoc comparisons. Error bars represent the standard deviation of biological replicates. Statistical significance is indicated as adjusted *P* < 0.05 (*), < 0.01 (**), and < 0.001 (***).

We subsequently assessed the ability of these mutants to form biofilm through crystal violet staining. Notably, only the *yjbH::tn* mutant demonstrated stronger biofilm formation during the time-course assay, with the most significant differences observed at 6 and 8 hours. Conversely, although inactivation of *mntR* diminished fibronectin adhesion, it did not exhibit a clear biofilm defect over the duration of the assay. However, a more thorough examination at 8 hours revealed a significant reduction in biofilm formation compared to the wild type (Fig. 4C). Complementation of *mntR* restored biofilm formation to wild-type levels at 8 hours, whereas complementation of *yjbH* resulted in biofilm formation below wild-type levels (Fig. 4C).

### *yjbH* and *mntR* serve as central regulatory hubs for biofilm formation across multiple lineages

Since *yjbH* and *mntR* are conserved across multiple *S. aureus* strains and appear to act as central regulators of biofilm-related functions, we wondered whether they would share functions across lineages. We investigated their effects in an alternative *S. aureus* strain, the methicillin-sensitive *S. aureus* (MSSA) strain SH1000 (ST8) (20). Surprisingly, a *yjbH* mutation in SH1000 increased adhesion to fibronectin to a greater extent that in JE2, but it did not affect its ability to form biofilm (Fig. 5). The effect of the *mntR* mutation on biofilm was also observed in SH1000 to a similar degree to that in JE2, but it did not exhibit a fibronectin adhesion defect (Fig. 5). Complementation of *yjbH* slightly increased biofilm formation in the mutant; however, it did not impact its ability to adhere to fibronectin. Complementation of *mntR* did not reverse the biofilm deficit caused by the *mntR::tn* mutant, possibly because our complementation plasmid used the JE2 sequence, and there may be small differences in regulation between JE2 and SH1000.

**Figure 5.**
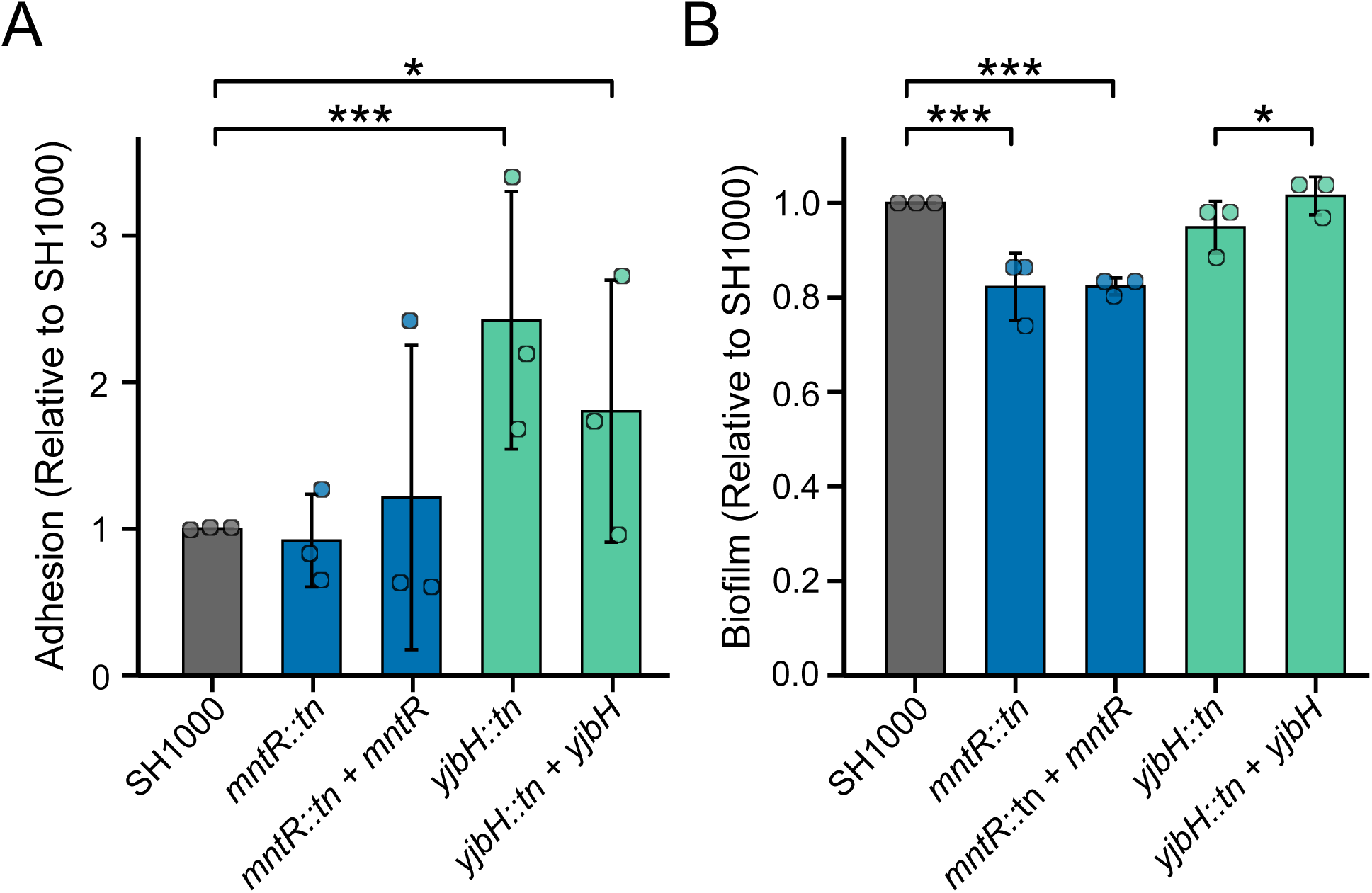
Genetic background influences the contribution of *yjbH* and *mntR* to *fnbA*-dependent adhesion and biofilm formation. (A) Fibronectin adhesion was measured in the SH1000 (MSSA) background. (B) Biofilm formation after 8 h of growth in TSBg was quantified by crystal violet staining. Data are presented as normalized adhesion (A) or normalized biofilm formation (B) relative to the WT. Error bars represent the standard deviation of biological replicates. Data were analyzed using linear mixed-effects models followed by Type III ANOVA and Dunnett-adjusted post hoc comparisons. Statistical significance is indicated as adjusted *P* < 0.05 (*), < 0.01 (**), and < 0.001 (***).

## Discussion

In many bacterial species, including *B. subtilis* (21), *Vibrio cholerae* (22), *Salmonella enterica* (23), and *Listeria monocytogenes* (24), biofilm formation is controlled by a central master regulator. Using this framework, we sought to identify potential master regulators that control two major components of the *S. aureus* USA300 biofilm, namely MSCRAMMs and eDNA. To do so, we harnessed the rapid, precise response of bioreporters to track biofilm dynamics across the NTML, achieving an unprecedented scale of bioreporter-based screening.

Our screen revealed that only a handful of mutants similarly affected both biofilm bioreporters, and none identified a clear, concise signalling pathway, suggesting the absence of a single central regulator coordinating these two matrix components. Notably, the matrix components investigated here fulfill distinct biological functions. They are expressed at different stages of growth (Fig. 1), which may inherently limit the existence of a unified regulatory control, since a regulator would need to orchestrate temporally and functionally diverse processes. A similar decentralized or stepwise organization of biofilm regulation has been described in *Pseudomonas aeruginosa*, where biofilm development relies on an integrated regulatory network rather than a single master switch. In this organism, biofilm formation is controlled at multiple levels, including quorum sensing (LasR-LasI), second messengers such as c-di-GMP, and a two-component system (GacA-GacS), whose interplay ensures a finely tuned and adaptive developmental program (25).

Although no shared central pathway emerged from both libraries, the pathways identified within each dataset independently were biologically coherent and consistent with previously described phenotypes. Genes involved in energy metabolism were significantly enriched among mutants with increased P*_fnbA_* activity (Fig. 3A). These mutants are typically associated with a small colony variant (SCV) phenotype, characterized by enhanced biofilm formation, increased adhesion, elevated internalization, and improved intracellular persistence within macrophages (6, 26), all of which have been associated with increased expression of adhesins such as FnbA.

Conversely, mutants with reduced P*_cidA_* activity were enriched for genes involved in riboflavin biosynthesis and flavin-containing metabolic pathways (Fig. 3B). Although a direct regulatory role for flavins or riboflavin in *cidA* expression is unlikely, flavin derivatives serve as essential cofactors in cellular redox reactions and are therefore critical for maintaining redox homeostasis (27). Alterations in flavin metabolism may consequently affect cellular redox status, thereby influencing CidA-mediated autolysis (28) and extracellular DNA release during biofilm development.

In *S. aureus*, redox balance is tightly regulated by multiple systems, including the Agr quorum-sensing pathway and the major redox-responsive regulator Spx, which orchestrates adaptation to oxidative stress (29). Several genes identified in both libraries are directly or indirectly linked to redox homeostasis, suggesting that oxidative stress management is an important regulatory axis influencing biofilm-associated processes (29). Notably, the shared hits *yjbH* and *mntR* highlight the contribution of distinct redox-related pathways to biofilm regulation. YjbH regulates ClpXP-mediated degradation of Spx, thereby modulating the oxidative stress response (29). In contrast, MntR is a manganese-dependent transcriptional regulator whose inactivation leads to constitutive expression of the Mn^2+^ importer MntABC, resulting in elevated intracellular manganese levels (30). Such disruption of metal homeostasis can impair proper metalation of proteins (31) and has been associated with increased susceptibility to oxidative stress in both *S. aureus* (32, 33) and *Escherichia coli* (34).

The oxidative burst is a major antimicrobial strategy of innate immune cells, particularly macrophages and neutrophils. It is characterized by the rapid production of reactive oxygen species (ROS) upon pathogen recognition. Although this response primarily occurs within the phagosome, ROS can also be generated extracellularly by assembling NADPH oxidase at the plasma membrane (35). ROS also promote the formation of neutrophil extracellular traps (NETs), which help contain *S. aureus* infections (36). Consequently, oxidative stress likely represents an important environmental signal encountered by *S. aureus* during infection. Consistent with this idea, ROS can damage iron-sulphur cluster-containing enzymes of the tricarboxylic acid (TCA) cycle, impairing respiration and promoting SCV-like phenotypes (37). Because SCVs exhibit enhanced biofilm formation and increased expression of surface adhesins, oxidative stress may promote biofilm-associated adaptations. However, further mechanistic studies will be required to determine whether the redox-associated regulator identified here contributes to this response.

Using this high-throughput bioreporter platform, we systematically analyzed the contributions of nearly 2,000 genes to the regulation of two major components of the *S. aureus* biofilm matrix. More broadly, this work demonstrates how dual-genetic reporters can transform genome-wide mutant libraries from static genetic resources into dynamic tools for probing the regulatory architecture underlying complex bacterial phenotypes. By simultaneously monitoring multiple pathways in real time, this approach captures regulatory interactions that conventional endpoint screens miss. Although applied here to biofilm development, the same strategy can readily be extended to virtually any genetically encoded phenotype, including stress adaptation, host-pathogen interaction, antimicrobial responses, or polymicrobial competition. We anticipate that this platform will facilitate the systematic dissection of complex bacterial behaviours under increasingly physiologically relevant conditions.

## Methods

### Strains and media

The bacterial strains used in this study are listed in Table S2. Most *S. aureus* strains were derived from MRSA USA300 JE2 (derived from the strain LAC isolated from the Los Angeles County jail in California), MSSA SH1000 (derived from NCTC8325-4), or RN4220 for genetic manipulation. *S. aureus* RN4220 was provided by François Malouin’s lab (Université de Sherbrooke). The *Escherichia coli* strain used for routine cloning was NEB 5 (New England Biolabs). For maintenance and preparation, *S. aureus* was grown from a single colony and incubated overnight with shaking (250 rpm) at 37 °C in Tryptic Soy Broth (TSB; 17 g/L casein peptone [pancreatic], 3 g/L soya peptone, 5 g/L NaCl, 2.5 g/L K_2_HPO_4_, 2.5 g/L glucose). For the biofilm assay, *S. aureus* was grown in TSB supplemented with 0.5% (w/v) glucose (TSBg). When needed, the following antibiotics were added to the media: chloramphenicol (10 µg/mL), erythromycin (5 µg/mL), and ampicillin (100 µg/mL), unless otherwise specified. Solid media were prepared with 1.5% (w/v) agar.

### DNA manipulation and plasmid construction

Plasmids used in this study are listed in Table S3. The pXen1 plasmid and ϕ85 phages were kindly provided by Eric Skaar’s lab (Vanderbilt University). P*_fnbA_*-*luxABCDE* and P*_cidA_*-*luxABCDE* reporters were constructed by amplifying the promoter regions using P787-P788 (P*_fnbA_*) or P803-P804 (P*_cidA_*) (Table S4), respectively, and inserting them into pXen1 between the EcoRI and BamHI restriction sites. Plasmids were transferred into *S. aureus* RN4220 by electroporation, then into JE2 by transduction (see below).

Assembly of the complementation plasmids was achieved by cloning into P*_fnbA_*-*luxABCDE.* Complementations, composed of the gene and its native promoter (500 pb upstream), were cloned in the opposite orientation relative to the promoter (P*_fnbA_*), using EcoRI with P956-P957 (*mntR*), or DraII with P969-P970 (*yjbH*). Sequences were verified to confirm that no other transcripts passed through the promoter and that the reporter itself is not affected. All plasmids were sequenced to confirm the absence of mutations.

### Reporter library preparation

All *S. aureus* strains were obtained by transferring constructs into *S. aureus* mutants from the Nebraska Transposon Mutant Library (NTML) via ϕ85-mediated generalized transduction (38). Phage amplification was performed by infecting 10 mL of RN4220 containing the plasmid at an OD_600_ of 0.2 with 250 µL of undiluted phage ϕ85 (∼3.75 x 10^9^ PFU) in a 1:1 TSB: phage buffer (50 mM Tris-HCl (pH 7.8), 1 mM MgSO_4_, 4 mM CaCl_2_, 90 mM NaCl, 0.1% (w/v) gelatin). After overnight infection, phages were harvested and separated from the remaining bacteria by filtration through a 0.22 µm filter.

High-throughput transduction was performed as described previously, with minor modifications (38). Briefly, the NTML mutants were grown overnight in 175 µL of TSB in 96-well plates, then infected with 20 µL of undiluted phage (∼3 × 10^8^ PFU) and 2.2 µL of 1 M CaCl_2_. After 30 minutes, 6.7 µL of 1 M sodium citrate was added to stop the infection, and the mixture was incubated for 1 hour. 7 µL was plated on TSA containing 5 µg/mL erythromycin, 20 µg/mL chloramphenicol, and 5 mM sodium citrate, and incubated overnight at 37 °C. A single isolated colony was then used to inoculate a well in a 96-well plate containing 150 µL of TSB with 5 µg/mL erythromycin, 10 µg/mL chloramphenicol, and 5 mM sodium citrate, and the plate was incubated overnight. Two subsequent passages were performed by transferring bacteria using a 96-pin replicator and growing them under the same conditions to eliminate all remaining phages. The libraries were frozen in 5 mM sodium citrate to prevent phage contamination. Mutants were randomly selected to confirm the presence of the reporter by light emission on a solid medium and the absence of phage contamination.

### High-throughput biofilm reporter assay

Manipulations of the reporter libraries were performed using a Rotor HDA (Singer instrument, United Kingdom). Individual library plates (384-well) were thawed for 1 hour and inoculated onto TSA containing 5 µg/mL erythromycin, 10 µg/mL chloramphenicol, and 5 mM sodium citrate. After 24 h of growth, colonies were transferred to 75 µL of TSB containing 10 µg/mL chloramphenicol and incubated for an additional 24 h. Mutants were then transferred into a white, optically clear polymer-bottom 384-well plate (Thermofisher, Cat# 142762) containing 50 µL of TSBg with 10 µg/mL chloramphenicol. Plates were immediately transferred to a pre-heated plate reader (TECAN Spark) at 37 °C. Reporter activity (Photons s^-1^) and bacterial density (OD_600_) were monitored every 10 min for 16 h without shaking.

### Screen analysis

Raw data were collected and analyzed using an R workflow. Briefly, OD_600_ was adjusted to an initial density of 0 by subtracting the minimum of the first 5 reads from each value. Reporter activity (Photons s^-1^) values were adjusted by subtracting a “blank” (mean of the first 5 hours) from each value. These adjustments reduced background values and allowed for a stronger analysis. Due to the relative expression formula (Photons s^-1^/OD_600_), low OD_600_ values (early time points) resulted in a large background. To avoid this, data with OD_600_ below 0.1 were excluded from the analysis. Peak activity was defined as the three highest successive values (the highest sum of 3 consecutive values), and the mean of these three was considered the peak activity. The OD_600_ for the first of these 3 values was considered the OD_600_ for the activity peak. Peak activity and its associated OD_600_ were normalized within each plate using IQM normalization (39). The distance between each value and the mean was then calculated using the z-score approach. The z-score is the deviation between the mutant’s actual value and the mean of the whole data, using the standard error. Mutants with |z-score| ≥ 1.96 were considered to deviate significantly from the population.

### Gene Ontology analysis

Gene Ontology (GO) enrichment analysis was performed using the BioCyc Pathway Tools Omics Dashboard to identify biological processes overrepresented among hits identified in the reporter screens. Because the JE2 genome is not comprehensively annotated in BioCyc, the analysis was performed using orthologous genes from the NCTC8325 genome. Orthologs were assigned using the aureowiki Gene ortholog list (https://aureowiki.med.uni-greifswald.de). Gene lists corresponding to mutants exhibiting increased or decreased reporter activity were analyzed separately. Statistical enrichment was assessed using a Fisher’s exact test with the *S. aureus* NCTC8325 genome as a reference. GO terms with a *P* < 0.05 were considered significantly enriched. Redundant, overlapping, or synonymous GO terms were manually removed to simplify data presentation.

### Biofilm formation and crystal violet staining

*S. aureus* biofilms were commonly prepared by diluting an overnight culture in 200 µL of TSBg in a 48-well plate to an initial OD_600_ of 0.005, corresponding to a bacterial density of approximately 1.75 x 10^5^ CFU/mL. Biofilms were grown statically for 24 h unless otherwise specified. The supernatant was discarded, and the biofilm was washed twice with 1X PBS, stained with 0.01% (w/v) crystal violet for 20 min, and then washed twice with water. Crystal violet was solubilized with 33% (v/v) acetic acid, and the optical density at 590 nm was measured using a plate reader (Biotek Synergy H1).

### Fibronectin adhesion assay

Wells in a 96-well plate were coated with 100 µL of fibronectin at 10 µg/mL (1 µg/well) overnight at 4 °C. They were washed once with PBS 1X and blocked with 200 µL of bovine serum albumin (BSA) at 2 mg/mL (400 µg/well) for 2 h at room temperature. Bacteria from an overnight culture were diluted to an OD_600_ of 0.05 in fresh TSBg and incubated for 2.5 h. Bacteria were then diluted to an OD_600_ of 0.4 (∼1.5 x 10^7^ CFU/mL) in PBS 1X, and 100 µL of this suspension was added to each well, which was then incubated at 37 °C for 90 minutes. Adhered bacteria were washed 3 times with PBS 1X, stained for 20 min with 0.1% (w/v) crystal violet, and washed 3 times with PBS 1X. Crystal violet was solubilized with 33% (v/v) acetic acid, and absorbance at 590 nm was measured using a plate reader (Biotek Synergy H1).

### Statistical analysis

Statistical analyses were performed using R version 4.5.2 (R Core Team, Vienna, Austria). All tests were performed using base R and the additional specified packages below. Depending on the experimental design, three complementary approaches were used, as detailed below.

#### Model-based analysis (biofilm dynamics)

The effects of condition and timepoint on biofilm were evaluated using a linear model (ordinary least squares regression; lm, *stats* package), including main effects and interaction (Condition x Timepoint). The overall significance of fixed effects was assessed using Type III analysis of variance (ANOVA; Anova, *car* package). Post hoc comparisons were performed using estimated marginal means (emmeans, *emmeans* package). Comparisons between each condition and the control were conducted at each time point using Dunnett-adjusted contrasts (contrast, method = “trt.vs.ctrl”, adjustment = “dunnett”) (Fig 4B).

#### Group-based analysis (single time point, independent wells)

Data distribution was assessed for each experimental group using the Shapiro-Wilk normality test (shapiro.test, *stats* package). When groups were considered normally distributed (*P* > 0.05), global differences were assessed using a one-way analysis of variance (ANOVA; aov, *stats* package). When at least one group deviates from normality, a non-parametric Kruskal-Wallis test (kruskal.test, *stats* package) was applied instead. Post hoc analyses were selected according to the experimental objective. For comparisons against a control strain, Dunnett’s test (DunnettTest, *DescTools* package) was used following ANOVA, whereas Dunn’s test with Benjamini-Hochberg correction (dunnTest, *FSA* package) was used following Kruskal-Wallis analysis. For all pairwise comparisons between groups, Tukey’s HSD test (TukeyHSD, *stats* package) was applied after ANOVA, and pairwise Wilcoxon tests (pairwise.wilcox.test, *stats* package) with Benjamini-Hochberg correction were used for non-parametric data. For predefined comparisons, pairwise t-tests (pairwise, *stats* package) or Wilcoxon tests were applied depending on the distribution, with Benjamini-Hochberg adjustment for multiple comparisons.

#### Linear mixed model (nested technical and biological replicates)

Data were analyzed with a linear mixed model (restricted maximum likelihood; lmer, *lme4* package), treating condition (strains, mutant, or complementation) as a fixed effect and biological replicate as a random intercept to account for non-independence among technical wells within each replicate. Normality of model residuals was assessed with the Shapiro-Wilk test. The overall effect of condition was tested with a type III ANOVA using Satterthwaite’s approximation for degrees of freedom (anova, *lmerTest* package). Post hoc comparisons were performed using estimated marginal means (emmeans, *emmeans* package). Comparisons between conditions of interest were conducted using a Benjamini-Hochberg-adjusted contrast (contrast, adjust = “bh”).

#### Global statement

Statistical significance was defined as *P* < 0.05. All statistical tests are reported in the corresponding figure legend.

## Acknowledgements

We thank members of the Beauregard, Côté, and Jean-Pierre laboratories for helpful discussions. We also thank Daniel Garneau for technical advice, Eric Skaar’s laboratory for providing pXen1 and phage Φ85, and François Malouin’s laboratory for providing strains. This work was supported by doctoral fellowships from the Fondation de l’Université de Sherbrooke (Bourse Hydro-Québec) and from the Fond de Recherche Nature et Technologie (https://doi.org/10.69777/315012) to J.S.B. and by a NOVA (NSERC and FRQ-NT) to P.B.B and J.P.C. (https://doi.org/10.69777/359578) as well as supporting funds from CR-CHUS to P.B.B and J.P.C.

## Author contributions

J.S.B. and P.B.B. conceived the project. P.B.B. supervised the development of the project. J.S.B. acquired data. J.S.B. and E.G. analyzed the data. J.P.C. and P.B.B. advised on data analysis. J.S.B. and P.B.B. wrote the manuscript, and all authors contributed to editing and approved the manuscript.

